# An accessible and generalizable in vitro luminescence assay for detecting GPCR activation

**DOI:** 10.1101/2023.03.02.530839

**Authors:** Ruby M. Miller, Jennifer Sescil, Marina C. Sarcinella, Ryan C. Bailey, Wenjing Wang

## Abstract

G-protein coupled receptors (GPCRs) serve critical physiological roles as the most abundant family of receptors. Here we describe the design of a generalizable and accessible **I**n vitro **G**PCR split **N**anoLuc l**i**gand **T**riggered **R**eporter (**IGNiTR**), having broad and diverse applications. IGNiTR leverages the interaction between a conformation-specific binder and agonist-activated GPCR to reconstitute a split nanoluciferase. We have demonstrated IGNiTR with three G_s_-coupled GPCRs and a G_i_-coupled GPCR with three classes of conformation-specific binders: nanobodies, miniG proteins, and G-protein peptidomimetics. IGNiTR demonstrated binding efficacy and potency values of various Dopamine Receptor D1 (DRD1) ligands that agree well with reported values. IGNiTR also allows the use of a synthetic G protein peptidomimetic, providing easily standardized reagents for characterizing GPCRs and ligands. We demonstrated three applications of IGNiTR: 1) characterizing GPCR functionality during Nanodisc-based reconstitution process; 2) highthroughput screening of ligands against DRD1; 3) detection of opioids for in the field applications. Due to its convenience, accessibility and consistency, IGNiTR will find extensive applications in GPCR ligand detection, screening and GPCR characterization.

## INTRODUCTION

G-protein coupled receptors (GPCRs) are a class of seven transmembrane proteins that function as essential intracellular signal transducers.^1^ Approximately 30% of FDA approved therapeutics target GPCRs.^2,3^ Due to the complexity of their signaling pathways and high modularity, GPCRs remain crucial targets for new therapeutic development.^4^ Live cell-based assays have been instrumental for GPCR drug screening, as well as GPCR signaling and mechanistic studies.^5-14^ However, there is still a lack of accessible and generalizable methods for detecting GPCR activation for broad applications. For example, it is infeasible with live cell assays to validate extracted GPCR protein’s functionality for biochemical and structural studies.

We envision an in vitro assay, based on GPCR protein in cell lysate, would provide a simple and easily adaptable format for broad applications. Existing in vitro assays, including radioligand binding, monitor GPCR-ligand binding, but do not measure ligand efficacy for inducing the active conformation, which recruits downstream G-proteins.^5,15-20^ An alternative approach, reconstitution of split bioluminescent enzymes, has been used to report on protein-protein interactions. Here we harness this robust luminescent signal that is quantifiable in a complex biological environment to track a ligand-induced binding interaction.^21-23^

We report the design of a highly adaptable GPCR luminescent assay for use in cell lysates and in solution. **IGNiTR** (**I**n vitro **G**PCR split **N**anoLuc l**i**gand **T**riggered **R**eporter) utilizes the agonist-dependent GPCR conformational change and subsequent recruitment of Gproteins and other conformation-specific binders^24,25^ to reconstitute split nanoluciferase (NanoLuc)^21,26,27^ (**Figure 1A**). Unlike live cell assays, IGNiTR components are easily stored with cell pellets expressing GPCR components frozen to preserve the integral, native lipid environment. Additionally, IGNiTR allows the use of peptidomimetics as conformation-specific binders, broadening assay applications.

**Figure 1.**
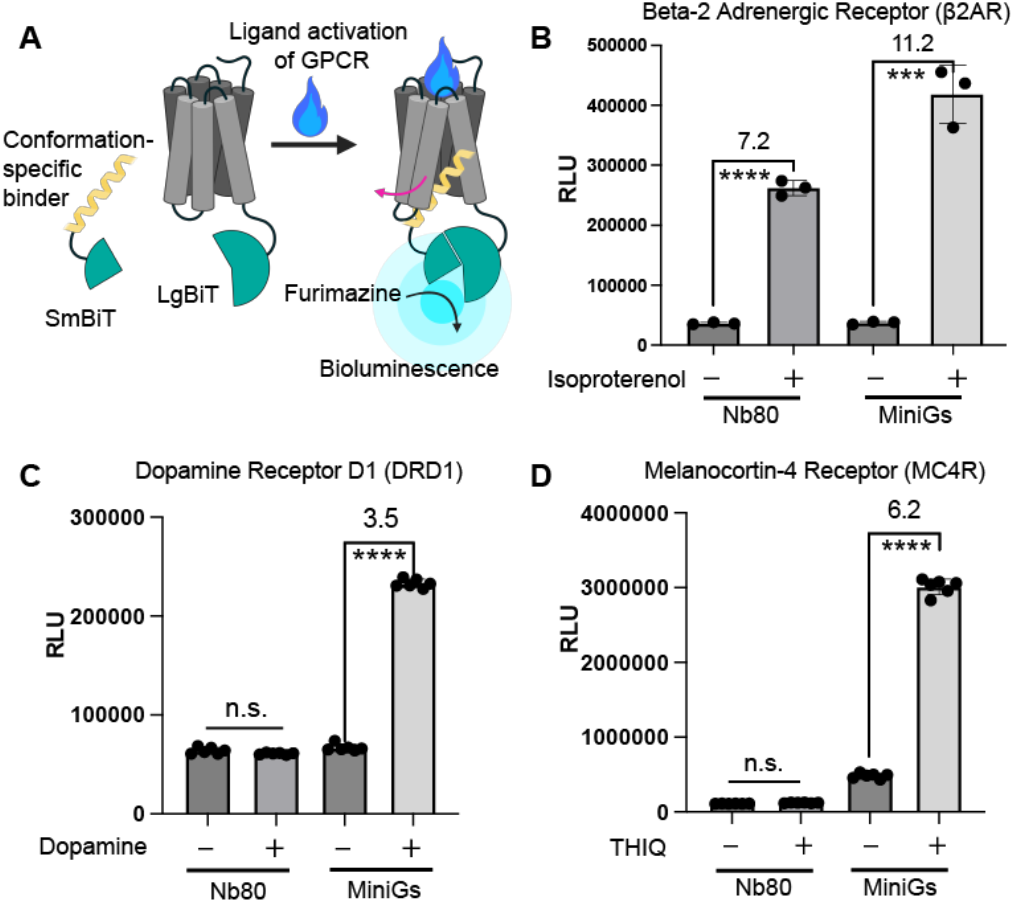
Characterization of IGNiTR with MiniGs and a conformation-specific nanobody. *A*. Schematics of the IGNiTR assay: LgBiT attached to a GPCR and SmBiT attached to a conformationspecific binder. Ligand activation of a GPCR results in GPCRconformation specific binder interaction, triggering split NanoLuc reconstitution. *B*. Characterization of IGNiTR with β2AR. *C*. DRD1, and *D*. MC4R with Nanobody 80 (Nb80) and MiniGs as the conformation specific binder. Drug, 10 µM. RLU: Relative Luminescent Units. n=3 for β2AR and n=6 for MC4R and DRD1. Values above the bars represent the DDR. Stars indicate significance after performing an unpaired Student’s t-test. ***P ≤ 0.001, ****P ≤ 0.0001. “n.s.” indicates no significant difference between the two conditions.

We have demonstrated the versatility of IGNiTR through three applications. First, IGNiTR proved a robust platform for high-throughput screening of ligands against Dopamine Receptor D1. Second, IGNiTR rapidly detects the µ-opioid receptor agonist fentanyl at nanomolar range in an easy and portable setup for potential field applications. Finally, IGNiTR was used to characterize protein functionality of GPCR protein samples at different stages of the Nanodisc-based GPCR extraction and reconstitution process. IGNiTR’s adaptability enables unique applications that are complementary to live cell assays or existing in vitro assays.

## RESULTS AND DISCUSSION

### Developing the IGNiTR assay

As shown in **Figure 1A**, IGNiTR is composed of two parts: the GPCR fused to one half of the split NanoLuc and a conformation specific binder fused to the other half of the enzyme. We began by using three different Gα_s_coupled GPCRs as models to develop IGNiTR: β2adrenergic Receptor (β2AR), Dopamine Receptor D1 (DRD1), and Melanocortin-4 Receptor (MC4R)^22^. For the conformation-specific binding components, we first explored the use of nanobodies and mini-G_s_ which have been used in live cell-based split NanoLuc assays.^21,23,28^ We tested Nanobody 80 (Nb80), which specifically binds activated β2AR.^29-32^ For a more generalizable application, we also tested miniG_s_ protein which has been shown to bind effectively to a range of the activated G_s_-coupled GPCRs.^28,30^

All IGNiTR constructs (prepared via sonication, see **SI** and **Figure S1**) produced significant ligand-dependent luminescence increase, hence a value > 1 for the ratio of the IGNiTR luminescence with drug to that without drug, abbreviated as drug dependent ratio (DDR) in this paper.

IGNiTR composed of β2AR fused to LgBiT and SmBiT fused to Nb80 or miniG_s_ each yielded significant DDRs (**Figure 1B**), indicating Nb80 and miniG_s_ can both selectively bind to the active conformation of β2AR in cell lysate. As expected, Nb80 did not show significant DDR with either DRD1 or MC4R. While when using miniG_s_, both DRD1 and MC4R produced significant DDRs (**Figure 1C & 1D**). The results validate that IGNiTR can detect a GPCR’s agonist-dependent conformational change in cell lysate. Furthermore, these results suggest that miniG_s_ is applicable for G_s_-coupled GPCRs.

### Using G-protein peptidomimetics in IGNiTR

To increase the versatility of IGNiTR, we used a Gprotein peptidomimetic as the conformation-specific binder in IGNiTR (**Figure S2**). Peptidomimetics enable the use of unnatural amino acids to increase binding synthesis facilitates the standardization of peptidomimetic concentration across large batches of well plates.

Our design was inspired by a reported G_s_ peptidomimetic^33^ (**SI** and **Figure S2**), which was based on the α-5-helix of Gα_s_ in the crystal structure of the G_s_ protein complex bound with β2AR.^35^ The G_s_ peptidomimetic (FNDCRDIIQRMHLRQYE-[Cha]-L) conserves the Gα_s_ protein α-5-helix amino acid sequence while adding a cyclohexylalanine (Cha) residue to increase hydrophobic interactions with the large hydrophobic pocket of the activated β2AR.^33,36^

Our design fused the SmBiT (11 amino acids) to the G_s_ peptidomimetic to create a SmBiT-G_s_ peptidomimetic fusion peptide. To test the peptidomimetic version of IGNiTR, β2AR-LgBiT protein in sonicated cell lysate was mixed with the G_s_ fusion peptide and NanoLuc substrate. Then, agonist or vehicle was added to evaluate the DDR (**Figure S3)**. β2AR IGNiTR with the G_s_ fusion peptide produced a significant DDR. We further tested the G_s_ fusion peptide with the other two G_s_-coupled GPCRs, MC4R and DRD1, each producing significant DDRs (**Figure 2A**). These results validated the G_s_ peptidomimetic’s selectivity for the active conformation of the G_s_-coupled GPCRs.

**Figure 2.**
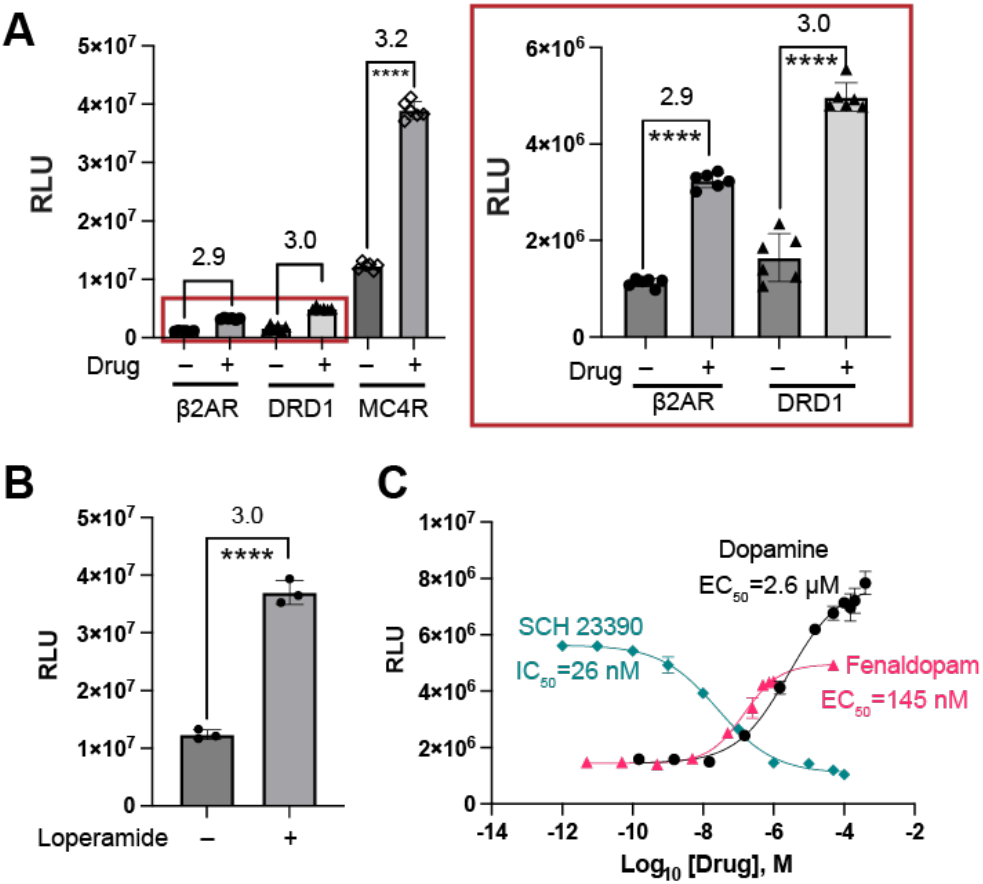
Characterization of the IGNiTR assay with peptidomimetics. *A*. Characterization of the IGNiTR assay with the Gs fusion peptide for β2AR, MC4R and DRD1. Gs fusion peptide, 2 µM; drug, 10 µM. n=6. *B*. Characterization of µ-OR IGNiTR with the Gi fusion peptide. n=3. Values above the bars represent the DDR. *C*. Dose-response curve of DRD1 IGNiTR with dopamine, fenaldopam and SCH 23390. For dopamine, EC50 range within 95% confidence is 1.8 to 4.1 µM. For fenoldopam, EC50 range within 95% confidence is 114 nM to 181 nM. For SCH 23390, IC50 range within 95% confidence is 20 nM to 33 nM. n=4. Stars indicate significance after performing an unpaired Student’s t-test. ****P ≤ 0.0001.

We next designed a G_i_ peptidomimetic using a similar strategy, since the α-5-helix of the Gα_i_-protein also interacts with a highly hydrophobic binding pocket based on the Gα_i_-protein structure.^37^ The G_i_ fusion peptide is composed of the SmBiT fused to the G_i_ peptidomimetic. We tested IGNiTR with the G_i_ fusion peptide and a G_i_coupled GPCR, the µ-opioid receptor (µ-OR). A significant DDR was observed for µ-OR IGNiTR (**Figure 2B)**. The result validates the G_i_ peptide’s selective binding to the affinity for the GPCR target.^33,34^ Additionally, peptide agonist-activated µ-OR and establishes the use of G_i_ fusion peptide in IGNiTR for G_i_ -coupled GPCRs.

### IGNiTR assay can characterize a GPCR ligand’s efficacy and potency

To further establish IGNiTR’s ability to characterize the various conformational states of a GPCR induced by various ligands, we applied the technique to DRD1 IGNiTR with full agonists, partial agonists and antagonists. The full agonist dopamine produced higher DDR than the partial agonist fenoldopam at saturated concentrations, with both producing a DDR > 1. The result validates that both full and partial agonists induce the active conformational state^38,39^ and that IGNiTR can differentiate ligand efficacies. DRD1 antagonist, SCH 23390, does not increase luminescence compared to the no drug condition.^38^ These results further validate the G_s_ peptidomimetic’s selective binding to the active conformation of DRD1. Lastly, DRD1 titration with dopamine and fenoldopam produced EC50 values of 2.6 µM and 145 nM, respectively (**Figure 2C**) that correspond well with the reported EC50 values.^38,39^ DRD1 titration with antagonist SCH 23390 in the presence of 10 µM agonist dopamine yielded an IC50 of 26 nM, which also agrees with the reported value.^40^ Overall, these characterizations demonstrate that the GPCR in IGNiTR maintains function comparable to live cell assays and IGNiTR can differentiate among various ligand efficacies.

### IGNiTR Application: High Throughput Screening (HTS) with robust Z’ values

IGNiTR could provide a valuable alternative for HTS of GPCR ligands, especially because IGNiTR components can be mixed in a batch, increasing consistency across large-scale screens. As a proof-of-concept, we performed a small-scale screen using DRD1 IGNiTR. We optimized DRD1 IGNiTR assay conditions by varying the DRD1 and G_s_ fusion peptide concentrations (**SI** and **Figures S4-6**). We then used the optimized DRD1-IGNiTR assay to scale up to screen for potential agonists using 1,916 compounds from an FDA-approved & Passed Phase I Drug Library from SelleckChem library. The Z’ value was consistent across the plates with an average of 0.79 (**Figure S7**) which is within the range of optimal Z’ value for HTS (1 > Z’ >0.5).^41^ Even though no hit molecules were identified from this library of compounds, this proof-of-principle screening demonstrated the feasibility and robustness of IGNiTR in GPCR ligand screening.

### IGNiTR Application: Rapid detection of opioids outside of a laboratory

IGNiTR could be packaged as an accessible kit for detecting GPCR agonists, such as opioids, outside of a biosafety level 2 laboratory space. The ongoing opioid crisis is being fueled by the emergence of additional synthetic opioids.^42,43^ There is a need for accessible methods to detect opioid derivatives which often are highly potent and thus have the potential to cause lethal overdoses.^43^ To address this need, we tested using μ-OR-based IGNiTR to detect the synthetic opioid, fentanyl.

First, we optimized the μ-OR and G_i_ fusion peptide concentrations for μ-OR IGNiTR (**Figure S6B, D)**. To increase the accessibility of IGNiTR for detection, we measured IGNiTR luminescence with a less sophisticated gel-imaging camera, rather than a plate reader. As shown in **Figure 3A & B**, higher concentrations of fentanyl result in increased luminescence intensity, with a plateau around 500 nM. μ-OR-based IGNiTR reliably detected 10 nM fentanyl which produced a significantly different intensity compared to the no drug control.

**Figure 3.**
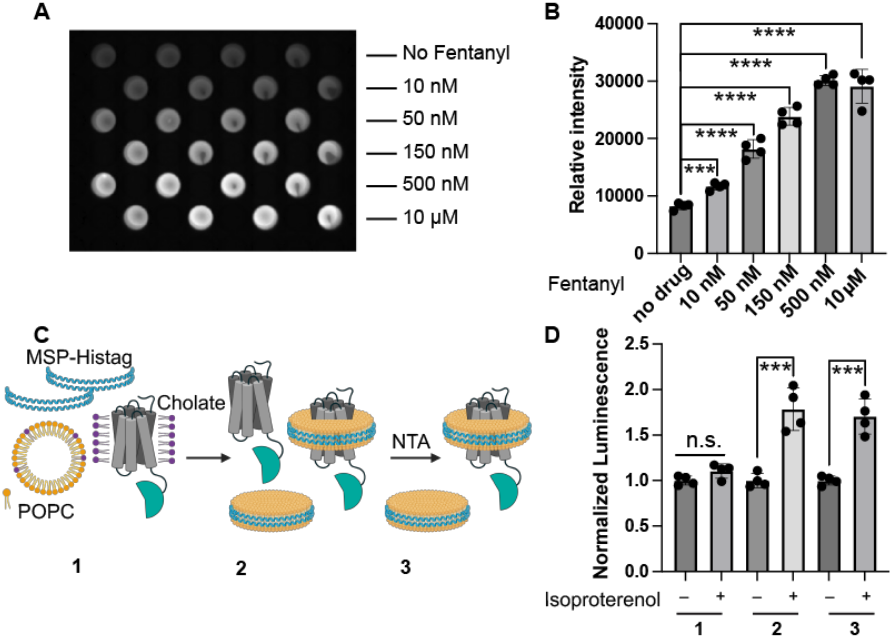
Applications of IGNiTR. *A. Imaging* and *B*. quantification of the IGNiTR assay performed with μ-OR LgBiT and the Gi fusion peptide to detect varied concentrations of fentanyl. n=4. *C*. Workflow for the incorporation of β2AR-LgBiT into POPC-based Nanodiscs. “NTA” represents Ni-NTA column purification. *D*. Analysis of the β2AR-LgBiT samples in C using IGNiTR with Gs fusion peptide. Stars indicate significance after performing an unpaired <u>Student’s t-test. ****P value<0.0001. ***P value<0.001.</u>

Our results demonstrate that IGNiTR can successfully detect various levels of opioids. Notably, IGNiTR reports on the general presence of opioids, which complements existing assays for detecting specific synthetic opioid molecules.^44,45^ Because the IGNiTR reagents can be readily stored frozen, we envision the components of IGNiTR being packaged into a kit for detecting μ-OR agonists in a variety of settings. Further, IGNiTR can potentially be adapted to detect other GPCR agonists, enabling biosensor development for a wide range of molecules.

### IGNiTR Application: Characterizing GPCR functionality during Nanodisc-based GPCR extraction and reconstitution

Nanodiscs have been widely applied for GPCR reconstitution by embedding the GPCR in a lipid bilayer, forming stable GPCR-lipid complexes.^46^ It’s particularly important but remains challenging to track the GPCR structural integrity and function throughout the Nanodisc assembly process.^47^ Therefore, we tested IGNiTR’s ability to characterize β2AR functionality during the three crucial steps of POPC-based Nanodisc formation^48^ as indicated in **Figure 3C**. Higher DDR suggests greater content of functional β2AR that can undergo agonist-dependent conformational changes, binding to the G_s_ peptidomimetic and reconstituting the split NanoLuc.

As shown in **Figure 3D**, β2AR reconstituted in Nanodisc with detergent cholate removed (sample 2) and its subsequent Ni-NTA purified sample (sample 3) produced a significant agonist-dependent DDRs, while the β2AR mixed with Nanodisc components as well as cholate (sample 1) did not yield a significant DDR. The result validates the importance of removing cholate for the correct folding and functionality of β2AR during its incorporation into the Nanodisc. The study establishes that IGNiTR could be used to monitor GPCR functionality throughout the protein extraction and reconstitution process, which is useful for optimizing these protocols.

## CONCLUSION

We have developed a generalizable in vitro GPCR assay, IGNiTR, that can characterize a GPCR’s structural integrity and activity by detecting the agonist-induced interaction of the GPCR with a conformation-specific binder. IGNiTR has features which can be both complementary to and advantageous over live cell-based assays. First, IGNiTR components, including the GPCR and the conformation-specific binder components can be prepared in advance and stored frozen until usage. Second, IGNiTR can be performed without the restrictions of working with live mammalian cells following biosafety level 2 regulations. Third, the preparation of IGNiTR in a cell lysate solution allows the use of a synthetic fusion G protein peptidomimetic, whose concentration can be well-controlled for assay fine-tuning, including optimization of DDR. Fourth and finally, mixing of the components can standardize the reaction conditions in thousands of wells to achieve consistency across HTS plates.

IGNiTR has advantages over existing in vitro assays. IGNiTR’s bioluminescent readout is quantifiable in a single step, and therefore can be easily scaled up and performed in the field while the existing in vitro GPCR assays, including the radioligand assay, requires a complicated set up.^19,20^ We demonstrated diverse applications in: 1) HTS of GPCR ligands; 2) characterization and detection of GPCR ligands in the lab and in the field; as well as 3) verifying GPCR structural integrity for in vitro GPCR characterizations. In future work, IGNiTR can be generally applied and expanded for all GPCRs.

## ASSOCIATED CONTENT

## Supporting Information

The Supporting Information is available free of charge on the ACS Publications website.

Supplementary figures and methods (PDF)

## AUTHOR INFORMATION

### Author Contributions

The manuscript was written through contributions of all authors. All authors have given approval to the final version of the manuscript.

## ACKNOWLEDGMENT

The research of the manuscript was funded by the University of Michigan, NIH MH132939. Ruby Miller is supported by NIH 1F31DA056192-01. We thank the Wang lab members, Dr. Shoji Maeda, Dr. Peter Toogood, Guanwei Zhou, Aubrey Putansu and Joseph Loomis for proof-reading the manuscript and providing edits. Figures created with BioRender.com.

