## Supplementary Information for "An accessible and generalizable in vitro luminescence assay for detecting GPCR activation"

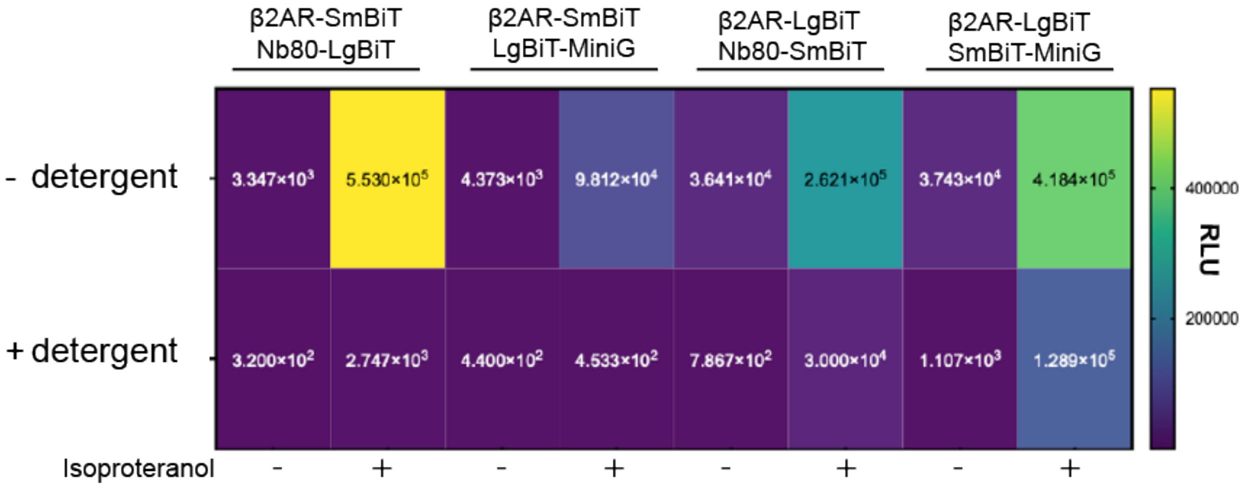


**Figure S1.** Comparison of IGNiTR component fusion geometry prepared in no detergent vs detergent conditions. Four versions of IGNiTR with different fusion geometries and Nb80 or miniG as conformation-specific binder were tested. The cell pellet expressing these IGNiTR components were lysed by sonication in solutions with or without the detergent mix (1% DDM (n-dodecyl β-D-maltoside) and 0.1% CHS (cholesteryl hydrogen succinate). Addition of detergent significantly decreases luminescence in all conditions. Therefore, we used sonication without detergent in the remaining work. Numbers inside the grids are relative luminescence signal.

A)

| | |
|---|---|
| Natural $G_s$ peptide | FNDCRDIIQRMHLRQYELL |
| SmBiT-$G_s$ peptide 1 | VTGYRLFEEIL-GS-<br>FN-C-SRD-C-IQRMHLRQYE-{Cha}-L |
| SmBiT-$G_s$ peptide 2 | VTGYRLFEEIL-<br>FNDCRDIIQRMHLRQYE-{Cha}-L |
| SmBiT-$G_i$ peptide | VTGYRLFEEIL-GSKKK-<br>FDAVTDIIIKENLKDCG-{Cha}-F |

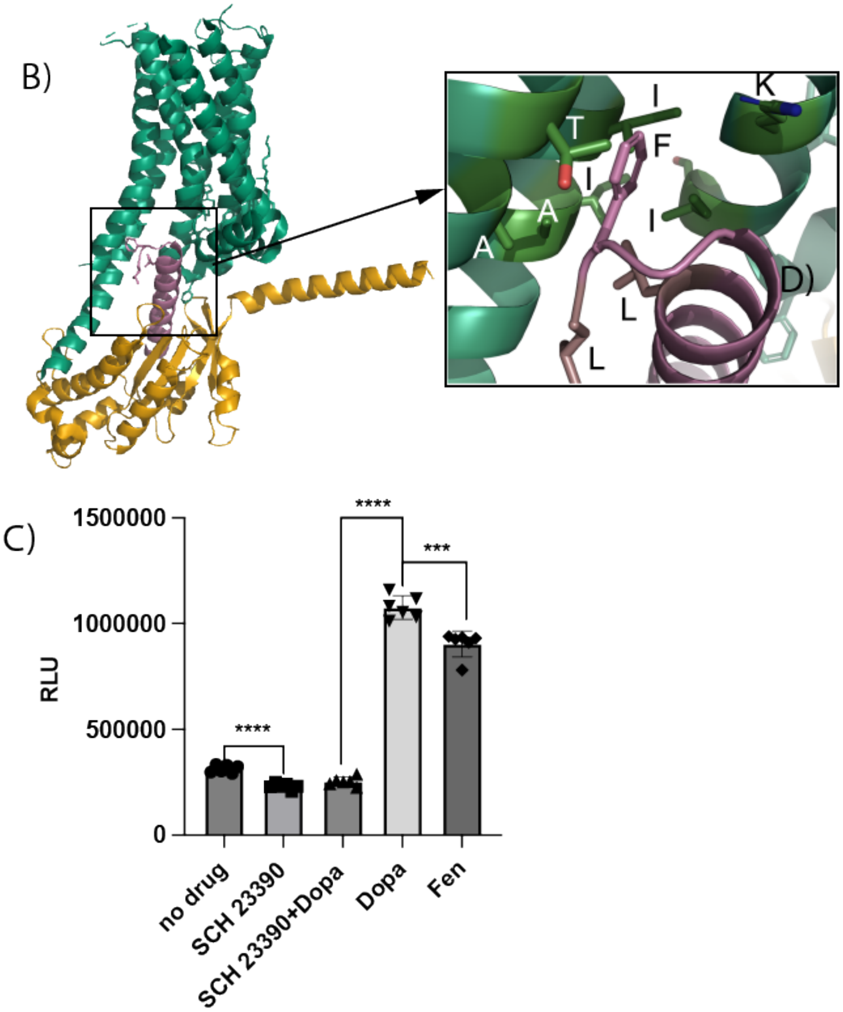


**Figure S2**. A. Amino acid sequences of the fusion peptides. *B.* Model structure based on LY3154207-bound DRD1 (PDB: 7X2F). Mutation phenylalanine is introduced in the penultimate position of the $G_s$ protein's $\alpha$-5-helix to illustrate the interaction between the $\alpha$-5-helix with the hydrophobic binding pocket of the activated DRD1. *C.* Comparison of DRD1-IGNiTR signal with a panel of drugs at saturated concentrations. SCH23390, 50 $\mu$M; SCH23390+Dopa (Dopamine), 50 and 10 $\mu$M; Dopa, 10 $\mu$M; Fen (Fenaldopam), 10 $\mu$M. Luminescence values were taken at 30 minutes post drug incubation.

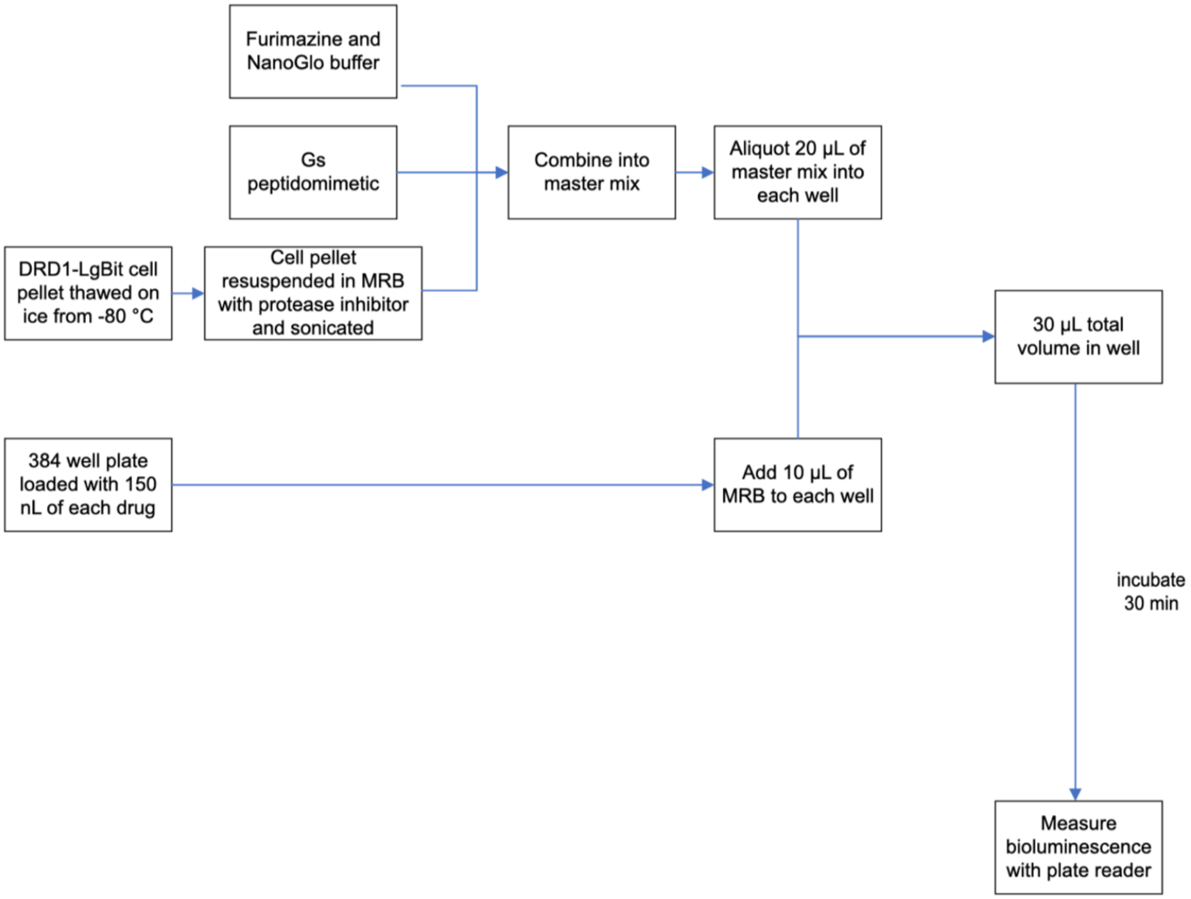


**Figure S3.** Flowchart showing preparation of IGNiTR assay. MRB is membrane resuspension buffer.

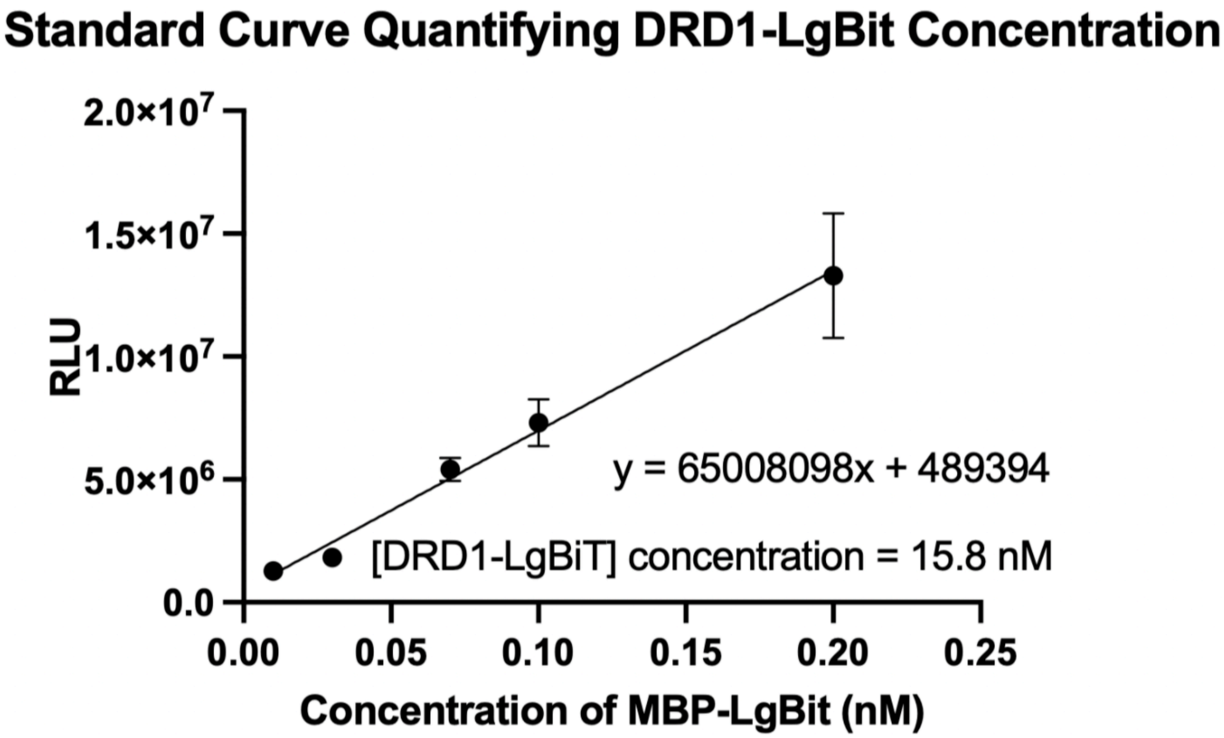


**Figure S4.** Standard curve using the luminescent signal of the high-affinity binder HiBiT with known concentrations of MBP-LgBiT to determine the concentration of DRD1-LgBiT.

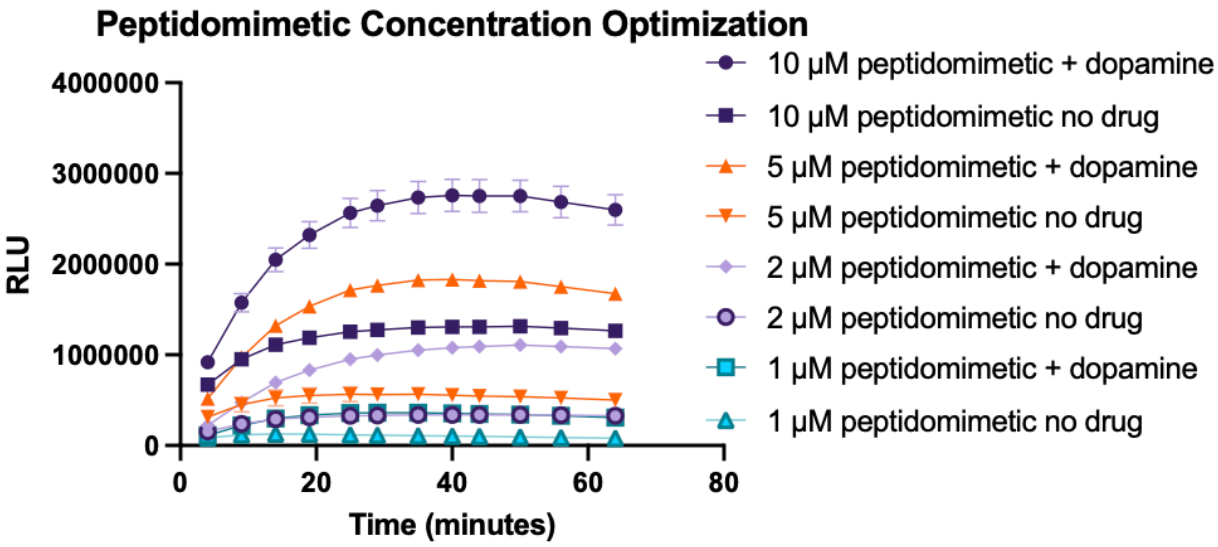


**Figure S5. Relevant to Figure 4B.** Time course of the luminescent signal of DRD1-LgBiT with different concentrations of peptidomimetics. Time 0, when drug is added. 10 μM dopamine was used.

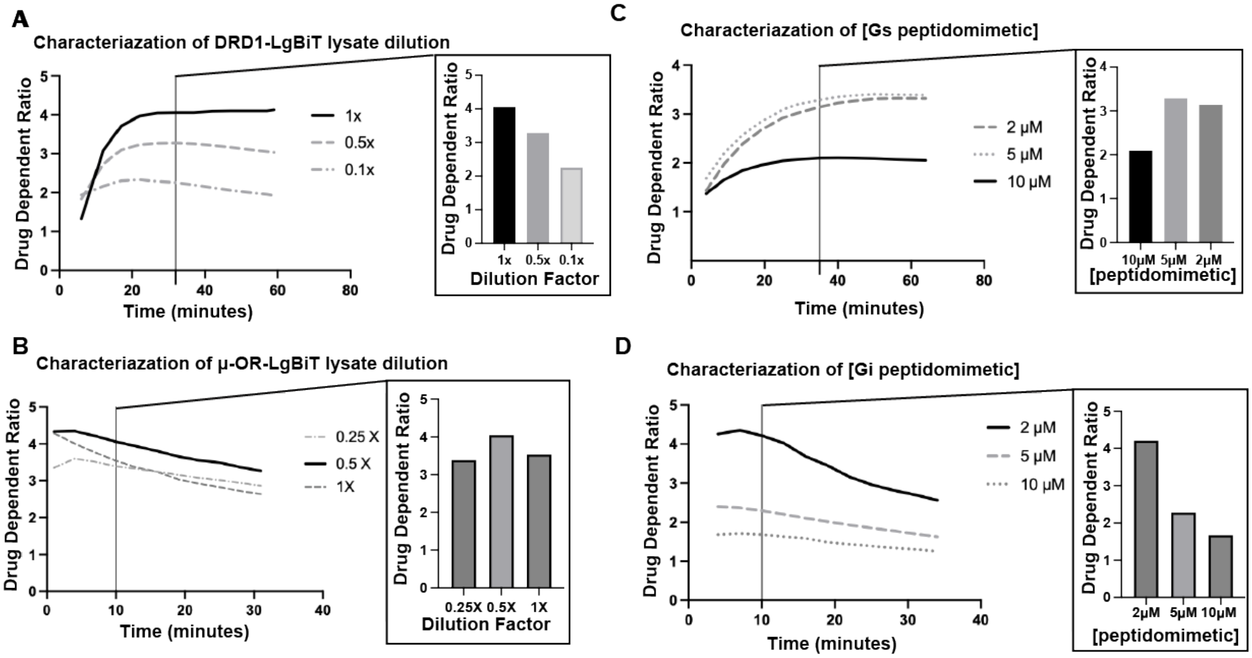


**Figure S6. Characterizing the IGNiTR assay with $G_s$ and $G_i$ fusion peptides.** *A.* Characterization of the effects of GPCR cell lysate dilution factors on IGNiTR using DRD1. *B*. Characterization of the μ-OR-based IGNiTR with cell lysate in a range of dilutions. *C.* Characterization of the effects of a range of $G_s$ peptidomimetic concentrations on IGNiTR using DRD1-LgBiT. DDR were taken around 30 minutes after drug incubation for both. N=6 *D*. Characterization of the effects of a range of $G_i$ peptidomimetic concentrations on IGNiTR using μ-OR-LgBiT. DDR were taken around 10 minutes after drug incubation for both. n=3.

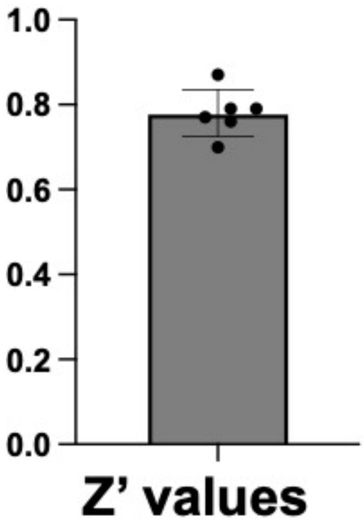


**Figure S7**. Z’ values measured across a proof-of-concept drug screening for DRD1-LgBiT and $G_s$ fusion peptide across 6x 384-well plates.

**Additional text:**

**Testing IGNiTR with varied geometry and lysis conditions with and without detergent**

We chose to express the GPCRs in mammalian cells to facilitate the correct folding, membrane trafficking and post-translational modification of these complex proteins.[1] Cell lysis is required for the G-protein mimic to access the intracellular loops of the GPCR and to enable storage of the cell pellet and ensure homogeneity of the lysate. Various methods have been developed for breaking up the cell membranes, as well as for solubilizing and stabilizing the GPCR protein, including the use of specific detergent mixes.[2] To maximally preserve GPCR protein folding and function, we tested two cell lysis conditions. We began by testing sonication of the cells in detergent-free solutions, since the native plasma membrane lipid environment provides crucial support for the structural integrity of membrane-bound GPCRs.[3] We also tested sonication in solutions containing a detergent mix frequently used to separate GPCR proteins from the membrane.

We tested two different fusion geometries with β2-adrenergic Receptor (β2AR) fused to either the large portion of the split NanoLuc, LgBiT, or the small portion, SmBit, and the G-protein mimic fused to the other half of the split NanoLuc. As shown in **Figure S1**, significant ligand-dependent luminescence increase was observed for both Nb80 and miniG$_s$ in the two geometries in the lysis condition without detergent.

The data in **Figure S1** also indicates that different fusions could affect IGNiTR performance. For example, when using Nb80 as the conformation-specific binder, much higher DDR was detected when GPCR was fused to SmBiT and Nb80 to LgBit. However, when using miniG$_s$ as the conformation specific binder, the opposite fusion geometry with GPCR fused to LgBiT and miniG$_s$ to SmBiT yielded a greater DDR.

**Figure S1** also shows that lysis with solutions containing detergent significantly diminished the luminescence in all conditions tested, suggesting the detergent disrupts the GPCR's functionality. The outcome was not unexpected since it has been shown that detergents can cause perturbations that affect the ability of the protein to be activated.[4] This experiment highlights the importance of keeping the GPCRs in their native lipid environment. Therefore, for optimal IGNiTR assay performance, the cell pellet will be lysed by sonication without detergent.

#### Testing $G_s$ peptidomimetics

Another version of the reported $G_s$ peptidomimetic incorporated two unnatural amino acids in the helical backbone away from the GPCR contact sites that form covalent bonds to generate stapled peptides to stabilize the helical structure. We decided to test both our non-stapled and this stapled $G_s$ peptidomimetic (**Figure S2**). However, for the stapled peptidomimetics, we tested using cysteine residues instead to facilitate a di-sulfide bridge formation in the α-helix.

We designed two fusion peptides composed of SmBiT fused to the two versions of $G_s$ peptidomimetics (**Figure S2**) and tested these fusion peptides in the IGNiTR assay using β2AR. While the disulfide-bond containing peptide 1 did not demonstrate agonist-dependence, $G_s$ peptidomimetic peptide 2 showed agonist-dependent DDR. The high background signal of $G_s$ peptide 1 can be attributed to the cysteine-based disulfide bond failing to connect the helical structure of the α-5-helix.

#### Finetuning of the IGNiTR assay with $G_s$ and $G_i$ fusion peptides

The advantage of an in vitro assay is that we could modulate the concentration of each of the components in the assay and fine tune the conditions. We therefore characterized the performance of the IGNiTR assay concerning following conditions: GPCR-LgBiT concentration, ligand incubation time, and $G_s$ fusion peptide concentration. To characterize IGNiTR with different GPCR-LgBiT concentrations, we first estimated the relative concentration of GPCR-LgBiT by creating a standard curve of LgBiT with its high-affinity peptide partner, HiBiT[5] (**Figure S4**). We then varied the dilution factor for the GPCR-LgBiT cell lysate while the peptidomimetic concentration was held constant at 10 μM. Among the dilutions, the 1x and 0.5x dilutions yielded the best DDR of >3 when measured at 30 minutes for DRD1 (**Figure S5 and 6**). Since the signal tends to stabilize around 25-30 minutes after ligand incubation, we therefore measured the luminescence at 30 minutes for all DRD1 characterizations. For the μ-OR-LgBiT, we observed that the DDR ratio peaks around 10 minutes, with all three dilutions, 0.25X, 0.5X and 1X, producing comparable DDR of ~ 4 (**Figure S6**).

We next characterized the IGNiTR assay with different concentrations of the fusion $G_s$ and $G_i$ fusion peptides. To minimize the cell pellet needed for characterization, we used the 0.5x of the GPCR-LgBiT lysate dilutions. We varied the concentration of fusion peptides added. For both $G_s$ and $G_i$ fusion peptides, the highest concentration (10 μM) resulted in a lower DDR compared to the lower concentrations due to higher background luminescence (**Figure S6**). Drug-dependent DDR ratios of ~ 3 were observed with 5 μM and 2 μM $G_{\alpha s}$ fusion peptidomimetic and the highest DDR of ~ 4 was observed with 2 μM $G_i$ peptidomimetic. We therefore used the 2 μM fusion peptide in the subsequent DRD1 and μ-OR IGNiTR assays to maximize DDR and to reduce the volume of fusion peptides needed for high throughput screening (HTS).

### Materials and Methods

#### *DNA Constructs and Cloning*

Standard cloning procedures, including NEB restriction enzyme digest, Q5 polymerase PCR amplification, T4 ligation and Gibson assembly, were used. Oligonucleotide primers were purchased from Sigma Aldrich. The plasmid DNA encoding MC4R was a gift from the Roger Cone Lab. The plasmid DNA encoding MiniGs and Nanobody 80 was purchased from Twist Biosciences.

Plasmid constructs were transformed into *Escherichia coli* cells via heat shock. XL1-Blue competent cells were used for all constructs, except for MBP-LgBit, which is described in "Expression and Purification of MBP-LgBit" below. Sequences were confirmed by Sanger Sequencing (Eurofins, GeneWiz)

#### *Cell Culture and Transfections*

HEK 293T/17 cell lines (ATCC, cat#: CRL-11268) used in these experiments were cultured at 37 °C and 5% $CO_2$. Cells were grown in complete growth media (1:1 MEM (Eagle's Minimal Essential Medium): DMEM (Dulbecco's Modified Eagle Medium, Gibco): with 50 mM HEPES (Gibco), 10% Fetal Bovine Serum (Sigma), and 1% Penicillin-Streptomycin (Gibco).

Cells at 80-90% confluence were plated into a flask that had been pre-incubated with human fibronectin for 10 minutes at 37 °C. An hour after seeding, these cells were transfected using FBS-free MEM and polyethlenimine (PEI, 1 mg/ml, Polysciences) with a ratio of 1:10 between µL of PEI and µg of plasmid DNA. The cells were then incubated at 37 °C for 20 – 24 hours.

The cell pellet was harvested by aspirating the media and resuspending the cells with a cell scraper in Dulbecco's Phosphate Buffered Saline (DPBS) buffer. 18 mL of DPBS was used to resuspend the cells within a T-75 flask, with this ratio kept constant for other flask sizes. 1.5 mL of resuspended cells were placed into an Eppendorf tube, which was centrifuged at 6,010 g for 3 minutes. The supernatant was aspirated, and the pellet was resuspended in DPBS, then centrifuged again under the same conditions. The supernatant was aspirated again, and the cell pellet was flash frozen in liquid $N_2$ and stored at -80 °C until ready for use.

*Preparation of Cell Pellet*

The cell pellets were prepared immediately before the assay, by first placing the cell pellet onto ice. For cell pellets containing GPCR constructs, the pellet was treated with 210 µL of Membrane Resuspension Buffer (MRB). MRB is comprised of incomplete membrane resuspension buffer (resuspension buffer comprising 20 mM HEPES, pH 7.5 and 2 mM $MgCl_2$) and benzoase (EMD Millipore, 70746) in a 4.5 mL to 1.8 µL ratio. For cell pellets containing cytosolic proteins (Nanobody 80 or MiniGs), the pellet was treated with Mammalian Protein Extraction Reagent buffer. After this, 2 µL of 100X protease inhibitor (Sigma Aldrich, P1860#) was added to a final concentration of 1X.and the pellet was resuspended by pipetting up and down. The solution was then sonicated using Model 50 Sonic Dismembrator (Fischer Brand) at a 20% amplitude (3 x 1 sec pulse) and returned to ice.

*IGNiTR Assay*

A master mix containing Nano-Glo Buffer and substrate (Promega, N2012), GPCR construct, and G protein mimic was prepared. For one well, the ratio was 9.75 µL of Nano-Glo Buffer, 0.25 µL of Nano-Glo substrate, 5 µL of GPCR pellet and 5 µL of the conformation specific binder. The conformation specific binder may be either peptidomimetic (LifeTein), miniGs pellet or Nanobody 80 pellet. The GPCR construct cell pellet was prepared separately and added to the master mix immediately prior to adding the master mix to the well. The concentrations were varied for the optimization assays (Figure 4), but the volumes remained constant. Concentrations for each experiment are indicated in the figure legends. The 384 well cell culture plates were preloaded with 10 µL of drug per well, followed by loading of the 20 µL of the master mix per well. Times reported on figure captions were recorded from the addition of the master mix to the first well of the plate. The luminescence values for all the conditions were measured using an EnVision 2104 Multilabel Reader (Perkin Elmer).

*High Throughput Screening (Supplementary Figure 7)*

To scale up the IGNiTR assay for high throughput screening, we used the Echo 655 (Labcyte) to load 150 nL of drug to each well of the 384 well plate. Then, the Multidrop Combi Reagent Dispenser (Thermo Scientific) was used to add 10 µL of MRB to each well. The same machine was used to add 20 µL of master mix, prepared as described in "*IGNiTR Assay"* using 2 µM fusion peptidomimetic and the 0.5x dilution of DRD1-LgBiT found to be optimal from the characterization in Figure 4. After a 30-minute incubation, the luminescence values were measured using an EnVision 2104 Multilabel Reader (Perkin Elmer).

$$Corrected\ Relative\ Luminescence = \frac{luminescence\ of\ sample - mean\ luminescence\ of\ negative\ control}{mean\ luminescence\ of\ positive\ control - mean\ luminescence\ of\ negative\ control} x\ 100\%$$

We validated the drug hits from an initial screen of six 384 well plates by testing four replicates of each compounds using the same method. From this validation, a dose response curve was constructed for promising candidates by loading 150 nL at a range of concentrations, using a mosquito X1 (SPT Labtech) to obtain final concentrations in well from 661 nM to 25.1 µM. 10 µL of MRB was loaded to the compounds, followed by 20 µL of master mix. After a 30 minute incubation, the luminescence values were measured using an EnVision 2104 Multilabel Reader (Perkin Elmer).

*Expression and Purification of MBP-LgBiT*

The DNA encoding MBP-LgBit was transformed into BL21 cells. A colony of these cells was inoculated in 5 mL Luria-Bertani broth with ampicillin at 37 °C overnight. The culture was then transferred to a 500 mL flask of Luria-Bertani broth with ampicillin and placed in a 37 °C shaker until OD-600 reached 0.4 to 0.8. Protein expression was induced by addition of 1000X 0.1g/mL IPTG, to a final concentration of 1X. The culture was then shaken overnight at room temperature.

The cells were centrifuged at 4,248 g for 5 minutes at 4 °C. The cell pellet was lysed by resuspension in cold Bacterial Protein Extraction Reagent (B-PER, Fisher) buffer with 1 mM dithiothreitol (DTT) and 1X protease inhibitor (BioBasic, BS386). 15 mL of B-PER was used for every 500 mL of bacteria culture. 3-4 µL of benzoase was added to the cells, followed by a 5 min incubation on ice to ensure full cell lysis. The cells were then centrifuged at 16,994 g for 10 minutes at 4 °C.

50 mL of clear lysate was added to 2 mL of Ni-NTA resin slurry and incubated at 4 °C for 10 minutes. This mixture was then purified via an Ni-NTA column. The purity of MBP-LgBiT was then established using gel electrophoresis and a Coomassie stain analysis.

*Determining the Concentration of MBP-LgBiT*

The molar extinction coefficient of MBP-LgBiT was calculated using Expasy PratParam to be 89,270 $M^{-1}$ $cm^{-1}$. This value was then used in conjunction with Bier's Law to establish the concentration of the protein by the absorbance at $A_{280}$. The absorbance was re-calculated for each time MBP-LgBiT was used.

*Standard Curve (Supplementary Figure 2)*

The standard curve was created using the same 384 well plates. First, a master mix containing a ratio of 5 µL of 30 µM HiBiT, 0.125 µL furimazine and 4.875 µL NanoGlo Buffer was prepared. 10 µL of this master mix was added to each well. 5 µL of MBP-LgBit or GPCR-LgBit was added, in the dilution ratio indicated in the figure legends. Times reported on figure captions were recorded from the addition of the MBP-LgBit or GPCR-LgBit to the first well of the plate.

*Statistical Analysis*

All statistical analysis was performed using GraphPad Prism 9 software. This software was also used to construct plots. The sample size is indicated in figure legends (where n is the number of independent replicates). The mean and standard error of the mean were calculated for each condition. Two-sided Student's t-tests were used to evaluate the significance between data points. Z' values were calculated using the following equation:

$$Z' = 1 - \frac{3SD\ of\ positve\ control + 3SD\ of\ negative\ control}{|mean\ of\ positive\ control - mean\ of\ negative\ control|}$$

where SD is the standard deviation.

*Fentanyl detection using IGNiTR*

The frozen pellet expressing μ-OR-LgBiT was thawed, resuspended, and mixed with the $G_i$ fusion peptide and NanoLuc substrate. The reaction mix was aliquoted to separate wells of an opaque white 96-well plate. A range of concentrations of fentanyl were added to the wells and the plate was imaged using an Azure Biosystems c600 for chemiluminescence. The resulting images were analyzed using imageJ.

*Nanodisc Assembly and Purification*

The pellet (β2AR-LgBiT) was resuspended in 400 uL membrane resuspension buffer (MRB) and sonicated. The lysate was quantified using the Pierce BCA protein assay kit (Thermo Scientific). The protein concentration was calculated using an average molecular weight of 40kDa for membrane proteins. Nanodiscs were made as described previously (ref-see below). POPC (Avanti Polar Lipids) was dried under nitrogen and stored in a desiccator overnight. POPC was solubilized to 50 mM with 100 mM sodium cholate. Nanodiscs were assembled by adding MSP1E3D1 (Millipore Sigma) and lysate to the solubilized lipids up to a final volume of 350 µL in standard disc buffer (20 mM Tris, 100 mM NaCl, 0.5 mM EDTA, 0.01% NaN3) supplemented with sodium cholate to a final concentration of 20 mM. The final lysate concentration in the mixture was 10 uM, the MSP:lysate was 4:1, and the lipid:MSP was 90:1. The component mixture was incubated on an end over end mixer at 4℃ for 45 minutes. 150 mg of Amberlite XAD-2 beads (Millipore Sigma) were added, and the component mixture was incubated at 4℃ overnight before the beads were removed. The resulting Nanodiscs were then purified with Ni-NTA spin columns (NEB). The purified Nanodiscs were then exchanged into standard disc buffer with Bio-Spin P-6 Gel Columns (Bio-Rad) to remove imidazole.
